# Genomic characterization of viruses associated with the parasitoid *Anagyrus vladimiri* (Hymenoptera: Encyrtidae)

**DOI:** 10.1101/2022.07.15.500286

**Authors:** Yehuda Izraeli, David Lepetit, Shir Atias, Netta Mozes-Daube, Gal Wodowski, Oded Lachman, Neta Luria, Shimon Steinberg, Julien Varaldi, Einat Zchori-Fein, Elad Chiel

**Affiliations:** University of Haifa, Department of Evolutionary and Environmental Biology, Haifa, Israel; Laboratoire de Biométrie et Biologie Evolutive, Université Lyon 1, CNRS, Villeurbanne, France; Newe Ya’ar Research Center, Department of Entomology, ARO, Ramat Yishai, Israel; Volcani Research Center, Department of Plant Pathology and Weed Research, ARO, Rishon LeZion, Israel; BioBee Sde Eliyahu Ltd, Emek Hamaayanot, Israel; University of Haifa – Oranim, Department of Biology and Environment, Tivon, Israel

**Keywords:** biocontrol agent, dicistrovirus, endosymbiont, iflavirus, mealybug, reovirus, virome

## Abstract

The knowledge on symbiotic microorganisms of insects has increased in recent years, yet relatively little data is available on non-pathogenic viruses. Here we studied the virome of the parasitoid wasp *Anagyrus vladimiri* (Hymenoptera: Encyrtidae), a biocontrol agent of mealybugs. By high-throughput sequencing of viral nucleic acids, we revealed three novel viruses, belonging to the families *Reoviridae* (provisionally termed AnvRV [Anagyrus vladimiri reovirus]), *Iflaviridae* (AnvIFV) and *Dicistroviridae* (AnvDV). Phylogenetic analysis further classified the AnvRV in the genus *Idnoreovirus*, and the AnvDV in the genus *Triatovirus*. The genome of AnvRV is comprised of 10 distinct genomic segments ranging in length from 1.5 to 4.2 Kbp, but only two out of the 10 open reading frames (ORFs) have a known function. AnvIFV and AnvDV each have one polypeptide ORF, which is typical to iflaviruses but very un-common among dicistroviruses. AnvRV was found to be fixed in a mass-reared population of *A. vladimiri*, whereas it’s prevalence in field-collected wasps was ~15%. Similarly, the prevalence of AnvIFV and AnvDV were much higher in the mass rearing population than in the field population. Transmission electron micrographs of females’ ovaries revealed clusters and viroplasms of *Reovirus*-like particles in follicle cells. AnvRV was not detected in the mealybugs, suggesting that this virus is truly associated with the wasps. The possible effects of these viruses on *A. vladimiri*’s biology, and on biocontrol agents in general are discussed. Our findings identify RNA viruses as potential players involved in the multitrophic system of mealybugs, their parasitoids and other members of the holobiont.

**Importance:** Different biological control approaches use industrially mass-reared natural enemy insects to reduce damage of arthropod pests. Such mass-reared cultures may be positively and/or negatively affected by various microorganisms, including viruses. Yet, current knowledge on virus diversity, especially in arthropods, is limited. Here, we provide the first virome characterization of a member of the wasps family Encyrtidae - the member being the parasitoid *Anagyrus vladimiri* - a commercially important natural enemy of mealybug pests. We describe the genome of three previously unknown RNA viruses, co-inhabiting this parasitoid, and elaborate on the prevalence of those viruses in individual wasps of both mass-reared and environmental origins. Microscopy images suggest that at least one of the viruses is transmitted maternally via the ovaries. We discuss the genomic structure of the viruses and the possible relationship between those viruses and the *A. vladimiri* host, with implications on improvement of biocontrol of mealybug pests.

## Introduction

Numerous arthropod species host symbiotic microorganisms that are integral to their life history, ecology and evolution (Zchori-Fein and Bourtzis, 2011). Such microorganisms engage in a wide range of symbiotic relationships, from parasitism to obligate mutualism, and may affect various biological features of their hosts (Zchori-Fein and Bourtzis, 2011; Drew et al., 2021). While the diverse interactions of bacteria, and to a lesser extent of fungi, with insects have been extensively studied, viruses associated with insects are much less explored. This is mainly due to paucity of knowledge of a great fraction of virus diversity as attested by recent large-scale or more focused surveys that unravelled numerous new virus lineages (Shi et al., 2016; Webster et al., 2016; Wu et al., 2020).

Among insects, it appears that endoparasitoids have special relationships with viruses. Those insects develop within an arthropod host, ultimately causing its death (Godfray, H. J., 1994). Most parasitoids belong to the order Hymenoptera. Apart from pathogenic viruses, they often harbour inherited viruses that may affect their phenotype. For instance, some viruses may affect their behaviour, as observed in the solitary parasitoid *Leptopilina boulardi*. In that case, the virus forces infected females to superparasitize (i.e., laying eggs in already parasitized hosts), thus favouring its horizontal transmission within the superparasitized host (Varaldi et al., 2003, 2005). The ssRNA iflavirus DcPV is also injected by the parasitoid *Dinocampus coccinellae* into its ladybeetle host during oviposition. The virus then replicates in the beetle’s brain which may participate in turning the ladybeetle into a ‘zombie’ that guards the parasitoid cocoon from predation (Dheilly et al., 2015). Other true free-living viruses, such as the entomopoxvirus DlEPV found in some parasitoid wasps, may protect the wasp offspring from host immune reaction (Coffman et al., 2020). On the farther end of the pathogenic-mutualist axis, are some domesticated viruses that became integrated into the insect genome. The best-known example of this phenomenon is the case of polydnaviruses (PDVs) which are associated with wasps that parasitize caterpillars of butterflies and moths. The endogenized viral genes allow the production of viral-like particles which are injected together with the eggs inside the caterpillar host and suppress its immune system, hence allowing the successful development of the wasp larvae (Herniou et al., 2013). Evidently then, viruses have diverse and important effects on the phenotype and ecology of endoparasitoids, and there are likely many more virus-insect associations that are yet to be unveiled. In the current study the viruses associated with *Anagyrus vladimiri* Triapitsyn (Hymenoptera: Encyrtidae), a solitary endoparasitoid wasp attacking mealybugs (Hemiptera: Pseudococcidae), were identified and characterized. *Anagyrus vladimiri* is mass-reared for commercial use in biological control programs to manage two major global pest species, which attack plants in dozens of families: the citrus mealybug, *Planococcus citri* (Risso) and the vine mealybug *P. ficus* (Signoret) (Bugila et al., 2015). The female parasitoid lays an egg into the body of the mealybug host, which continues to develop while the wasp larva feeds and develops inside it. After a few days the mealybug host dies, its cuticle hardens and turns into a ‘mummy’ and the parasitoid larva pupates inside. Eventually, the adult parasitoid chews a hole in the mummified host and emerges (Avidov et al., 1967).

Recently, we reported an analysis of the fungal and bacterial portion of microbiome inhabiting this population of *A. vladimiri*. We found that the symbiont *Wolbachia* is dominating the bacterial community and under some conditions it has minor negative effects on the fecundity of the parasitoid (Izraeli et al., 2020). The current research was launched to characterize the virome of *A. vladimiri*, annotate the viral genomes, assess their phylogenetic position, and test the prevalence of the various viruses in lab and field populations of the wasp.

## Results

### *Electrophoresis characterization of* Anagyrus vladimiri *virome*

Viral nucleic acids (VNAs) were purified from a pool of adults (from the mass-reared line, see Table 1 for details) and treated by RNaseA and DNaseI. At least nine bands ranging in size from ~1.5Kbp to ~4Kbp were clearly observed when VNAs were separated on non-denaturating gel by electrophoresis (Fig. 1, lane 3). VNAs were successfully digested by RNaseA (Fig. 1, lanes 8-11), but not by DNaseI (Fig. 1, lanes 13), suggesting that the virome of *A. vladimiri* is composed of RNA viruses only. For both our positive control (Phi6 dsRNA) and the VNAs from *A. vladimiri*, the RNase digestion was mitigated in high concentration of sodium citrate buffer (=high ionic strength) with low concentration of RNase, which is the expected outcome when the RNA is double-, and not single-stranded (Sorrentino et al., 1980; Khramtsov et al., 1997). These results suggest that the tested strain of *A. vladimiri* harbours at least one, dsRNA virus with nine segments (or more, since segments of similar size may co-migrate at the same position), or several non-segmented or segmented dsRNA viruses.

**Fig 1.**
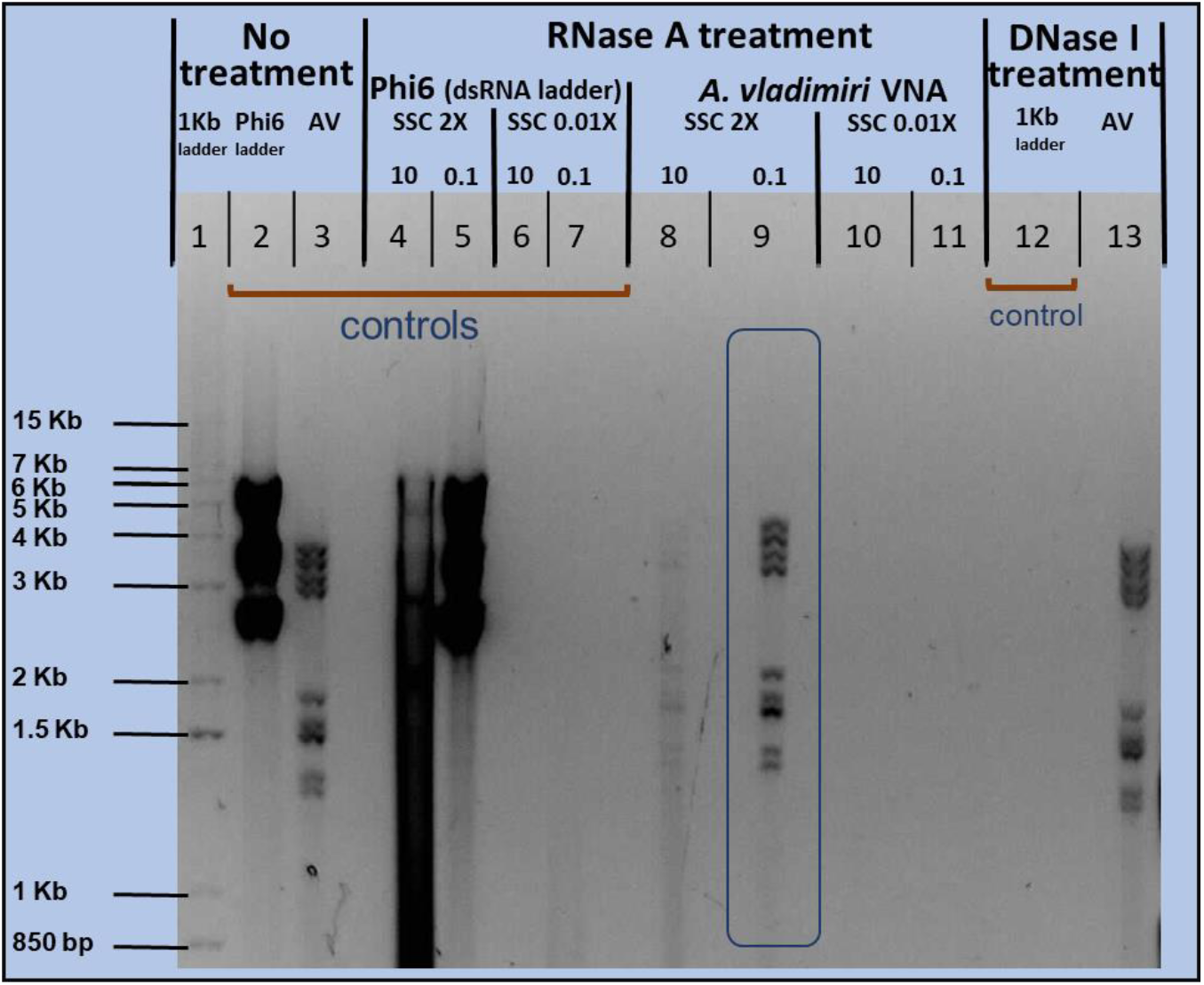
Electrophoresis pattern (using a 0.8% agarose gel) of VNAs extracted from *A. vladimiri*, which were treated by RNase and DNase separately. ‘AV’ denotes VNAs of *A. vladimiri*. ‘SSC’ denotes sodium citrate buffer, which was used in 0.01x and 2x ionic strengths. ‘10’/‘0.1’ denotes high/low concentration (μg/ml) of RNase respectively. Details by lane numbers: 1-3; no digestion treatment. 1; 1Kb ladder (1KB+, Invitrogen, MA, USA), 2; Phi6 – a ladder made of viral dsRNA (Invitrogen, MA, USA), 3; *A. vladimiri* VNA. 4-11; Digestion by RNase A in four buffer/enzyme concentrations. 4-7; Phi6 ladder as a control for the enzymatic activity of RNase. 8-11; AV digestion. 12-13; Digestion by DNase I, 12; 1Kb – a DNA ladder as a control for the enzymatic activity of DNase. 13; AV digestion.

**Table 1.** Summary of *Anagyrus vladimri* lines used in this study.

| **Table 1.** Summary of *Anagyrus vladimiri* lines used in this study. |  |  |
| --- | --- | --- |
| *line name* | *source* | *experiments used for* |
| **MR** | ‘Biobee’ mass-rearing facility | Virome analysis |
| **W<sup>+</sup>RV<sup>+</sup>** | ‘Biobee’ mass-rearing facility, and further reared in our lab.<br>Wasps carry *Wolbachia* and AnvRV. | Virus prevalence screening,<br>Reovirus segment screening,<br>Transmission electron microscopy |
| **W<sup>-</sup>RV<sup>+</sup>** | MR wasps that fed on antibiotics (Izraeli *et al.* 2020)<br>Wasps carry AnvRV but not *Wolbachia*. | Virus prevalence screening |
| **Field** | Field collections, north Israel | Virus prevalence screening,<br>Reovirus segment screening<br>(negative controls) |
| **Field-RV<sup>-</sup>** | Iso-female lines of the field collected wasps | Presence of viruses in mealybugs |

### *Sequencing the RNA virome of* A. vladimiri

After reverse transcription, the VNAs of *A. vladimiri* were sequenced on an Illumina platform. We obtained 16 million of high quality paired-end reads.High sequence duplication levels were observed (>96%), suggesting that the genome sequence is complete (except for the extremities of untranslated regions), and as expected from the apparent low complexity of the sample (Fig. 1) which totalled approximately 20-30Kbp. De novo assembly of this dataset, resulted in only 30 contigs, 18 of which were shorter than 1,500bp.

Homology (by amino acid sequence identity) of 11 out of the 12 contigs longer than 1,500bp, as identified by BLASTx and PSI-BLASTp, were found to three RNA virus families; *Dicistroviridae, Iflaviridae* (both positive strand ssRNA, order: *Picornavirales*) and *Reoviridae* (segmented linear dsRNA, order: *Reovirales*) (Table 2, Fig. 2).

**Fig 2.**
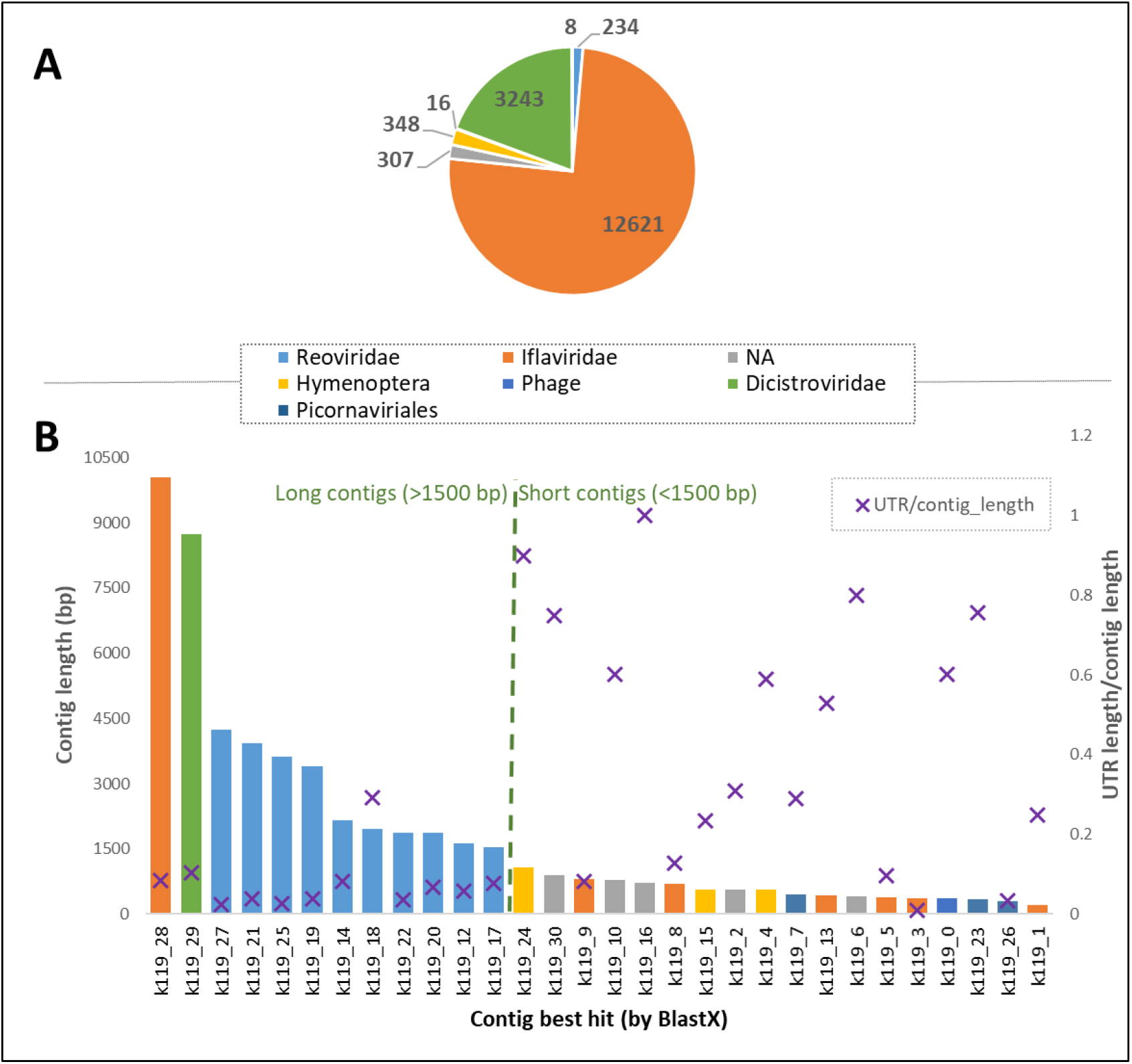
Virome analysis of *A. vladimiri*. The assembled 30 contigs were assigned to seven taxonomic groups according to the best BLASTx hit. **A)** Average depth per taxonomic group, calculated as: ((sum number of reads*100bp)/sum length of contigs). **B)** Columns represent the size of each of the 30 contigs (bp length). Purple crosses represent the proportion of both 5’ and 3’ untranslated regions (UTRs) to the contigs’ full length, as predicted by the ORF finder. High proportion suggests insufficient protein products, while low proportion suggests a true protein is translated from this contig. See table 2 for more details about each contig.

**Table 2.** Virome analysis of *A. vladimiri*. Details for all 30 contigs resulted from RNAseq.

| contig name | contig length | ORF length | UTRs length | UTR/contig length | BLASTx hit | taxonomic group | amino acid identity (%) |
| --- | --- | --- | --- | --- | --- | --- | --- |
| k119_28 | 10031 | 9173 | 858 | 0.085535 | YP_009111311.1 | Iflaviridae | 47 |
| k119_29 | 8726 | 7829 | 897 | 0.102796 | NP_620564.1 | Dicistroviridae | 38 |
| k119_27 | 4250 | 4148 | 102 | 0.024 | YP_392501.1 | Reoviridae | 33 |
| k119_21 | 3936 | 3788 | 148 | 0.037602 | QBA09478.1 | Reoviridae | 29 |
| k119_25 | 3620 | 3521 | 99 | 0.027348 | APG79179.1 | Reoviridae | 25 |
| k119_19 | 3388 | 3257 | 131 | 0.038666 | YP_392504.1 | Reoviridae | 45 |
| k119_14 | 2163 | 1985 | 178 | 0.082293 | YP_392505.1 | Reoviridae | 22 |
| k119_18 | 1963 | 1388 | 575 | 0.292919 |  | Reoviridae* |  |
| k119_22 | 1880 | 1814 | 66 | 0.035106 | QYV43128.1 | Reoviridae |  |
| k119_20 | 1865 | 1739 | 126 | 0.06756 | AWA82241.1 | Reoviridae | 24 |
| k119_12 | 1620 | 1526 | 94 | 0.058025 | YP_392507** | Reoviridae |  |
| k119_17 | 1535 | 1418 | 117 | 0.076221 | QBA09478.1 | Reoviridae | 34 |
| k119_24 | 1067 | 107 | 960 | 0.899719 | XP_017796731.1 | Hymenoptera |  |
| k119_30 | 887 | 223 | 664 | 0.748591 |  | NA |  |
| k119_9 | 809 | 743 | 66 | 0.081582 | YP_009342053.1 | Iflaviridae | 39 |
| k119_10 | 782 | 311 | 471 | 0.602302 |  | NA |  |
| k119_16 | 711 | 0 | 711 | 1 |  | NA |  |
| k119_8 | 701 | 611 | 90 | 0.128388 | YP_009342053.1 | Iflaviridae | 40 |
| k119_15 | 571 | 437 | 134 | 0.234676 | CAB0037418.1 | Hymenoptera |  |
| k119_2 | 564 | 389 | 175 | 0.310284 | XP_015596585.1 | NA |  |
| k119_4 | 560 | 230 | 330 | 0.589286 | EZA52908.1 | Hymenoptera |  |
| k119_7 | 451 | 320 | 131 | 0.290466 | CAA84692.1 | Phage |  |
| k119_13 | 432 | 203 | 229 | 0.530093 | YP_009342053.1 | Iflaviridae |  |
| k119_6 | 412 | 83 | 329 | 0.798544 |  | NA |  |
| k119_5 | 394 | 356 | 38 | 0.096447 | QFP98431.1 | Iflaviridae | 36 |
| k119_3 | 375 | 371 | 4 | 0.010667 | YP_009342053.1 | Iflaviridae |  |
| k119_0 | 358 | 143 | 215 | 0.600559 | ABD33943.1 | Picornavirales |  |
| k119_23 | 351 | 86 | 265 | 0.754986 | AGF91671.1 | Phage |  |
| k119_26 | 306 | 296 | 10 | 0.03268 | QGJ03578.1 | Phage |  |
| k119_1 | 218 | 164 | 54 | 0.247706 | YP_009342053.1 | Iflaviridae |  |

\*concluded as part of the genome of AnvRV by the segment screening assay (see table 3)
\*\*this hit was found only by using the PSI-BLASTp algorithm

One contig was most similar to a *Dicistroviridae* member named Black queen cell virus infecting honeybees, with an average of 41% amino acid sequence identity. The contig is 8,726bp long, which is in the expected range for *Dicistriviridae* in general (8.5-10.2kb) and very close to the genome length of Black queen cell virus (8,550bp). We propose to name this newly discovered virus: Anagyrus vladimiri dicistrovirus (AnvDV).

One contig, 10,031bp long, had the highest similarity to different members of the family *Iflaviridae*, with 46% amino acid identity with the iflavirus member *Dinocampus coccinellae* paralysis virus used as a reference. The contig length is well within the expected range for *Iflaviridae* (8.8-9.7kb). We propose to name this newly discovered virus: Anagyrus vladimiri iflavirus (AnvIFV).

Nine contigs were identified as homologous to nine different segments of a reovirus infecting the winter moth *Operophtera brumata* (ObIRV). Protein alignment to the known ObIRV segments as a reference by BLASTp, revealed that the coverage between the nine contigs to the nine segments ranges between 20-98%, while the amino acid sequence identity ranges between 19% and 37% in all segments (Table 2). The one remaining contig out of the 12 ‘long’ ones, had no similarity according to BLASTx and Psi-BLASTp searches (contig k199_18). However, its length was similar to other contigs assigned to the reovirus, therefore it was added to the screening assay for all reovirus segments in individuals. Results of that assay (see below) strongly support the assumption that this contig is part of the reovirus genome, and together with the nine contigs identified by BLAST, this genome consists of 10 segments. We propose to name this newly discovered segmented virus: Anagyrus vladimiri reovirus (AnvRV).

Six out of the 18 contigs shorter than 1,500bp (contigs k119_1, k119_3, k119_5, k119_8, k119_9 and k119_13; Fig. 2), were also assigned to the *Iflaviridae* family by the BLASTx search (Table 2, Fig. 2). Aligning those to the long AnvIFV contig showed that the small contigs are actually parts of this long contig with >96% nucleotide identity. They were discarded from the analysis.

The nearly-complete gnome sequence of the three novel RNA viruses is available in GeneBank under BioProject PRJNA641546, with the following BioSample accession numbers__.

### Annotation

Analysis of the recovered contigs predicted that all 12 ‘long’ (>1,500bp) ones have a large open reading frame (ORF), covering >90% of their length, suggesting that they encode true functional genes. In contrast, most of the 18 ‘short’ (<1,500bp) contigs had large portions of un-translated regions (UTRs), with the ORFs covering on average less than 60% (Table 2, Fig. 2).

Interestingly, the contig identified as AnvDV harbours only one ORF, unlike most known viruses of this family, which harbour two ORFs. Furthermore, no ‘internal ribosome entry site’ (IRES) - an RNA element typically found between the two ORFs of dicistroviruses (and enables the translation initiation of the second ORF) - was found along this contig. The genomes of the three RNA viruses were further annotated by searching the predicted ORFs for conserved domains using the NCBI CDD tool. Both the AnvIFV and AnvDV were found to each harbour five domains encoding for: RNA-dependent RNA polymerase (RdRP) (accession: cd01699), RNA helicase (accession: pfam00910), capsid proteins (CRPV_capsid superfamily, found in many dicistroviruses) (accession: cl07393) and two ‘rhv-like’ domains (part of a capsid protein, named as ‘drug-binding pocket’ found in poliovirus, also a member of *Picornavirales* order) (accession: cd00205) (Fig. 3a).

**Fig 3.**
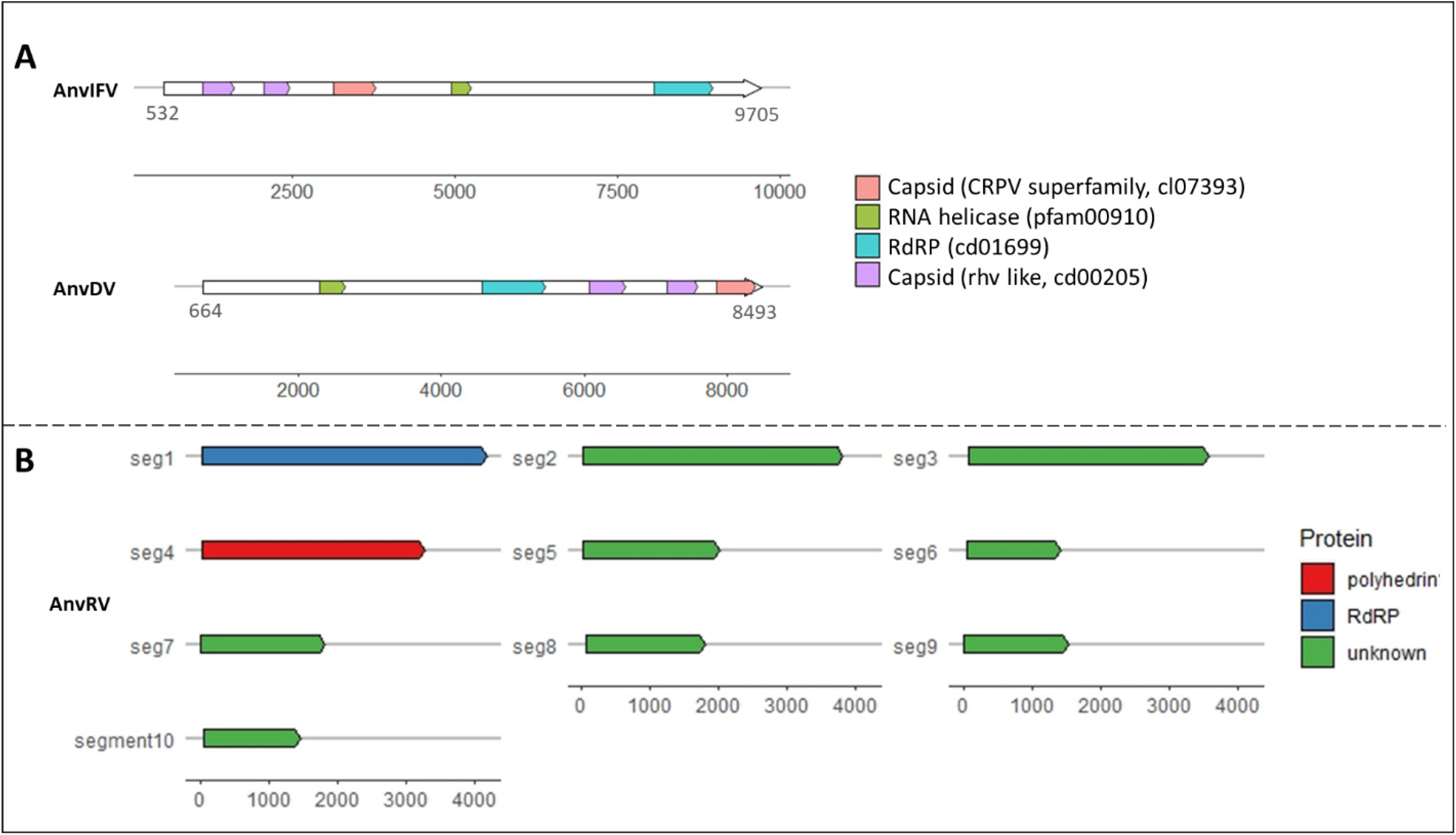
Genome annotation of the three novel RNA viruses. **A)** AnvDV and AnvIFV. White box indicates location of ORF, colored boxes indicate identified protein domains. Numbers on scale are in bp according to the full contig. RdRP, RNA-dependent RNA polymerase. **B)** Annotation of the segmented AnvRV. The ten segments are drawn to scale, with boxes indicating the ORF location on the full segment, colors represent the proteins encoded from those segments as assigned from homology to known sequences of the ObIRV genome.

No conserved domains were found for the 10 AnvRV segments. Nevertheless, two genes were inferred from the BLASTx searches by homology with the ObIRV genome: RdRP on segment no.1 and a polyhedrin protein on segment no.4 (Fig. 3b).

One conserved sequence motif (GAAGAKC) was found in the 3’ end termini of the five out of 10 contigs assigned to AnvRV (supp. fig 2). No conserved sequences were found in the 5’ end, suggesting that a few bases in the extremities of the contigs may be missing in our current assembly.

### Phylogeny

#### AnvRV

Phylogenetic analysis of the RdRP encoding segment of the AnvRV revealed that its closest known relative is the ‘Zoersel tick virus’, identified from the tick *Ixodes ricinus*. This virus, together with three other close relatives, have no genus taxonomic classification. The closest classified virus is the ObIRV, which belongs to the genus *Idnoreovirus* (Fig. 4). Pairwise alignments of the AnvRV to those five viruses reveals a 34-43% RdRP amino acid sequence identity, while it’s amino acid identity with all other viruses compared in the phylogenetic analysis was 27% and lower. As the common cut-off between ICTV verified genera is 30% amino acid identity (Matthijnssens et al., 2022), this suggests that the AnvRV, together with the four other unclassified relatives, belongs to the genus *Idnoreovirus* (Fig. 4).

**Fig 4.**
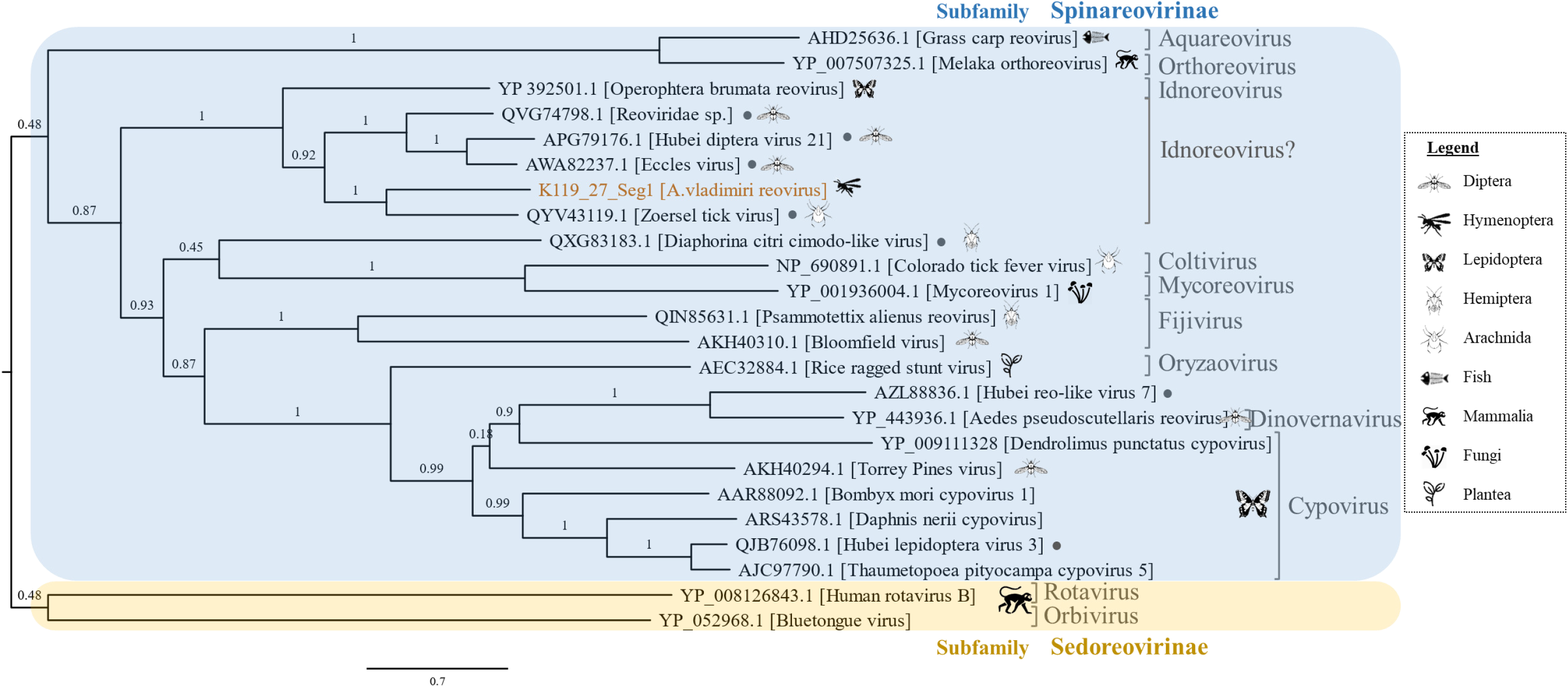
Phylogenetic analysis of AnvRV (colored), based on amino acid sequence similarities of the RdRP genome segment (~1,200aa long). Representative viruses of all nine genera in the subfamily Spinareovirinae are shown. The icons of organisms on the tips represent the taxonomic group of the host in which the virus was found. The • symbol indicate viruses that have not been classified to the genus level. The human Rotavirus B and the bovine Bluetongue virus (Family: *Reoviridae*, subfamily: *Sedoreovirinae*) were used as an out-group. The tree was inferred by maximum-likelihood using the LG substitution model, the numbers on branches indicate the results of approximate likelihood ratio tests for branch support (aLRT-SH test; Guindon et al. 2010). Scale bar: 0.7 substitutions per site.

#### AnvDV

The phylogenetic analysis for the AnvDV clearly show that this virus belongs to the genus *Triatovirus*, together with the Black queen cell virus (51% amino acid RdRP identity) and other relatives (Fig. 5).

**Fig 5.**
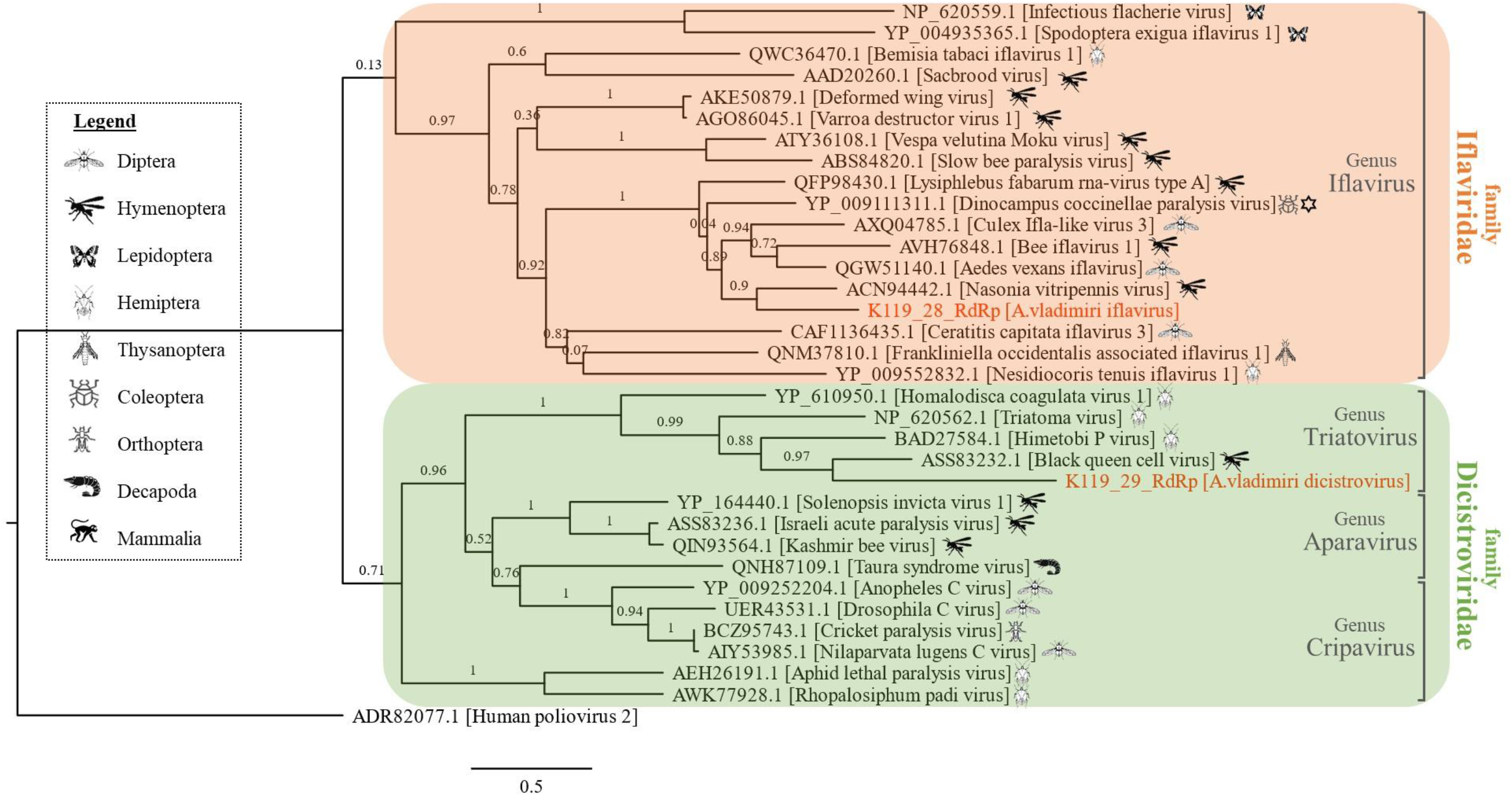
Phylogenetic analysis of A. vladimiri iflavirus (AnvIFV) and A. vladimiri dicistrovirus (AnvDV) (both colored), based on amino acid sequence similarities of the RdRP gene (~300aa long). The human poliovirus (order: *Picornavirales*, family: *Picornaviridae*) was used as an outgroup. The icons of organisms on the tips represent the taxonomic group of the host in which the virus was found (however, in some cases the viruses can be found in hosts from additional arthro-pod orders, such as Cricket paralysis virus). The ‘star of David’ symbol signs the virus that is reported to have a mutualistic relationship with its insect host. The phenotype of all other viruses is either unknown or pathogenic. The tree was inferred by maximum-likelihood using the LG substitution model, the numbers on branches indicate the results of approximate likelihood ratio tests for branch support (aLRT-SH test; Guindon et al. 2010). Scale bar: 0.5 substitutions per site.

#### AnvIFV

The phylogenetic analysis for the AnvIFV place this virus near the Nasonia vitripennis virus [accession ACN94442.1], Aedes vexans iflavirus [QGW51140.1](61-62% amino acid RdRP identity), and others. The unique Dinocampus coccinellae paralysis virus [YP_009111311.1], also clusters on the same branch with AnvIFV, and has 58% amino acid RdRP identity with it (Fig. 5).

### Prevalence of viruses in lab and field samples

To assess the prevalence of the three RNA virus candidates in the different *A. vladimiri* populations, diagnostic PCRs with specific primers were used (Supp. table 1) on 20 individual females from each of the *Wolbachia-carrying* (W^+^) and *Wolbachia-free* (W^-^) lab-reared lines (Izraeli et al., 2020). In addition, 24 field-collected individuals were screened (see Table 1 for details).

All 40 lab-reared individuals harboured the AnvRV. The AnvIFV and the AnvDV were found in 33 and 25 specimens out of 40 individuals respectively (Fig. 6). All three viruses were also detected in field individuals, although the prevalence was lower, with three females carrying AnvRV (12.5%), 10 carrying AnvDV (42%), and two carrying AnvIFV (8%) out of 24 individuals (Fig. 6).

**Fig 6.**
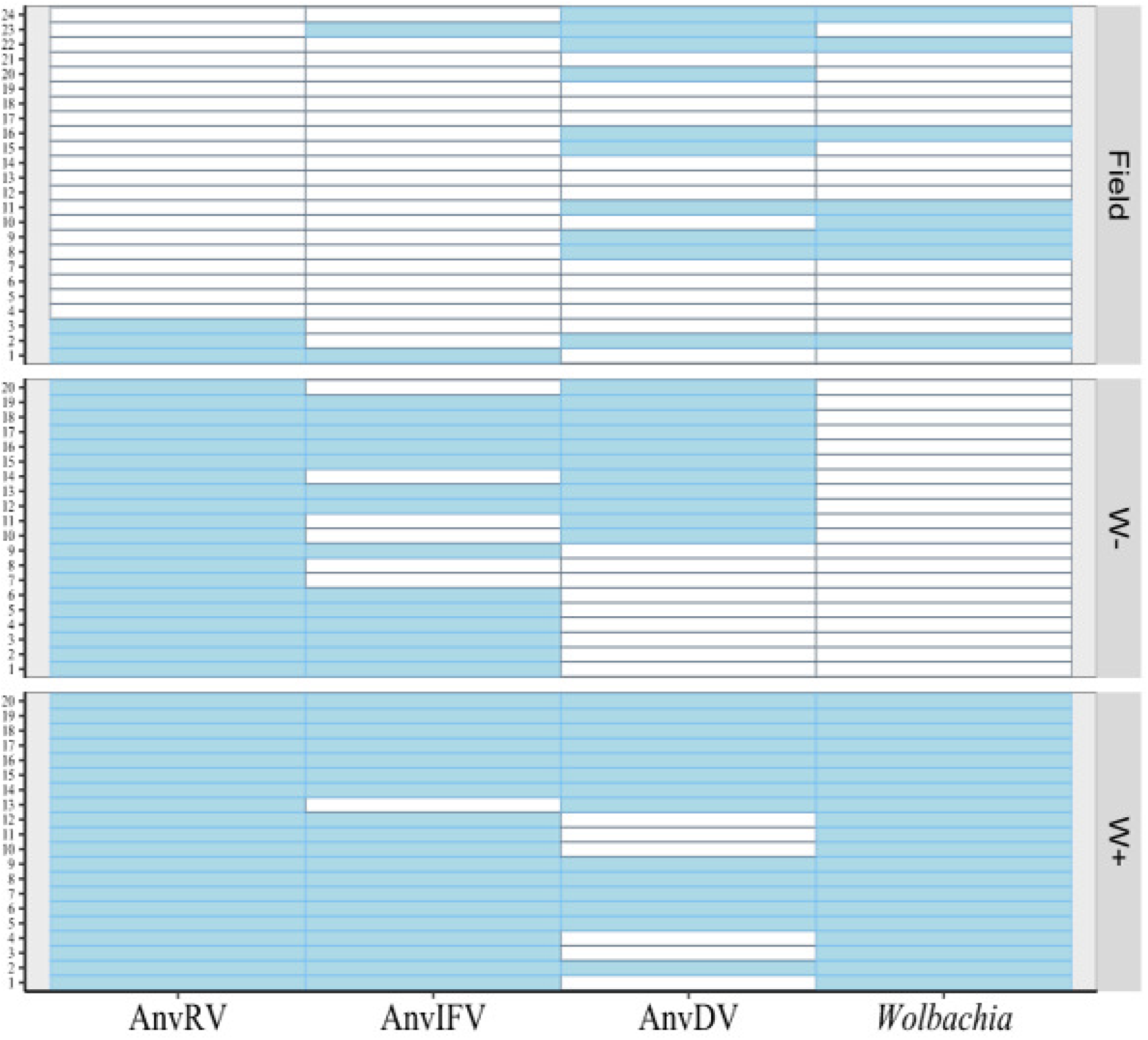
The multi-infection status of three lines of *A. vladimiri* individuals by the three RNA viruses and *Wolbachia*. Boxes on the same horizontal axis denote the infection status of the four microbes in the same individual; blue fill=detected, no fill=undetected. Detection of the AnvRV, which has a segmented genome, was done with the primers targeted to the largest segment, encoding for the RdRp gene (see supp. table 1 for all primer pairs used). W+: wasps from the *Wolbachia-infected* line, W-: wasps from *Wolbachia*-uninfected line.

Interestingly, out of 10 field females carrying the AnvDV, only three lacked *Wolbachia*, and seven out of the eight females carrying *Wolbachia*, also carried the AnvDV. The co-occurrence of those two microbes was statistically significantly correlated (Fisher’s exact test: p=0.002), suggesting a positive interaction between *Wolbachia* and AnvDV within *A. vladimiri* populations. However, AnvDV was present both in the *Wolbachia* positive and the *Wolbachia* negative strains in the lab.

### *Screening for the 10 Reovirus segments in individual* A. vladimiri *wasps*

To test whether all the 10 Reovirus segments (nine identified by sequence homology, and one - contig K119_18 - suspected by its length and UTR/contig length ratio) belong to the same AnvRV genome, rather than to different RNA viruses, 24 individuals from two wasp lines were screened by a set of diagnostic PCRs. Those included 16 individuals from the W^+^RV^+^ line - a line harbouring the AnvRV as confirmed by PCR with primers targeting the RdRP segment, and 8 individuals from the ‘Field-RV^-^’ line, as negative controls. Generally, the results were as expected (table 3); in 9 W^+^RV^+^ individuals all the segments were amplified, while in 6 ‘Field-RV^-^’ individuals, none were amplified, suggesting that all 10 segments belong to the same virus. However, in 7 individuals, one or two segments failed to amplify, and in two individuals from the negative population one single segment was detected (Table 3).

**Table 3.**
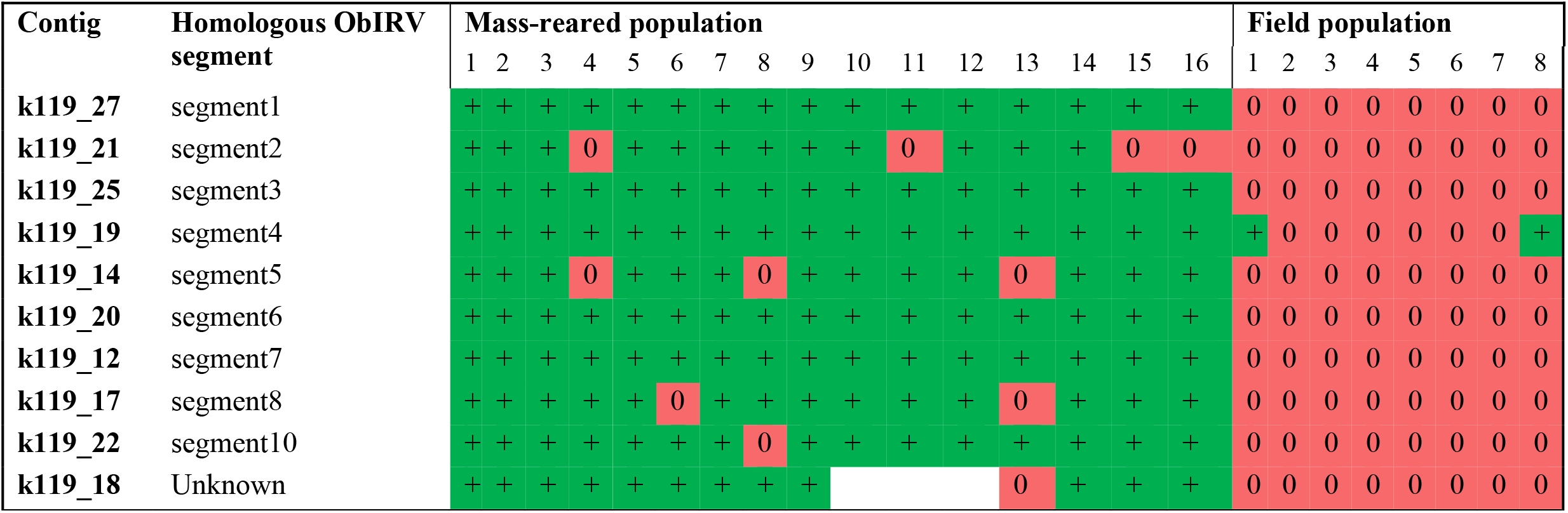
Screening of ten AnvRV segments in 24 individuals from two lines of *A. vladimiri*; 16 females from the MR population, and eight from Field population. ‘0’ denotes absence, ‘+’ denotes presence of the viral genomic segment by diagnostic PCR. Empty boxes denote untested samples.

### Presence of viruses in the mealybugs

To test whether any of the three RNA viruses originated from the mealybug hosts, four pools were tested for their presence: i) Mass-reared (MR) W^+^ *A. vladimiri*, ii) Field-collected *A. vladimiri* line without the AnvRV (RV^-^), iii) un-parasitized *P. citri*, and iv) *P. citri*, four days after parasitization by MR *A. vladimiri*, (see Table 1 for details). AnvRV was not detected in the un-parasitized mealybugs, nor in the Field-RV^-^ *A. vladimiri*, which were reared under the same conditions as the MR *A. vladimiri*, indicating that this virus was not acquired from the mealybugs, and is associated with *A. vladimiri*. In contrast, AnvIFV and AnvDV were detected in both *A. vladimiri* and un-parasitized mealybugs (albeit the bands for the mealybug samples were weak, they were confirmed by Sanger sequencing, with >99% identity), suggesting that those viruses may be acquired from the mealybugs or from the rearing environment, possibly the potato sprouts that the mealybugs feed on (Cheng et al., 2021), or even the various fungi that develop on this substrate (supplementary Fig. 1).

### Virus particles in A. vladimiri ovaries

To validate the presence of viruses in the females’ reproductive tissues, transmission electron microscopy (TEM) was carried on sections of *A. vladimiri* ovaries (from the W^+^RV^+^ line, see Table 1 for details). Images obtained from the three specimens tested showed clusters of round shaped viral particles, 60-65nm in diameter, organized either in viroplasm structures, or in dense clusters in the perinucleus area in the cytoplasm of follicle cells (Fig. 7). This pattern is typical to *Reoviridae* viruses (Shah et al., 2017).

**Fig 7.**
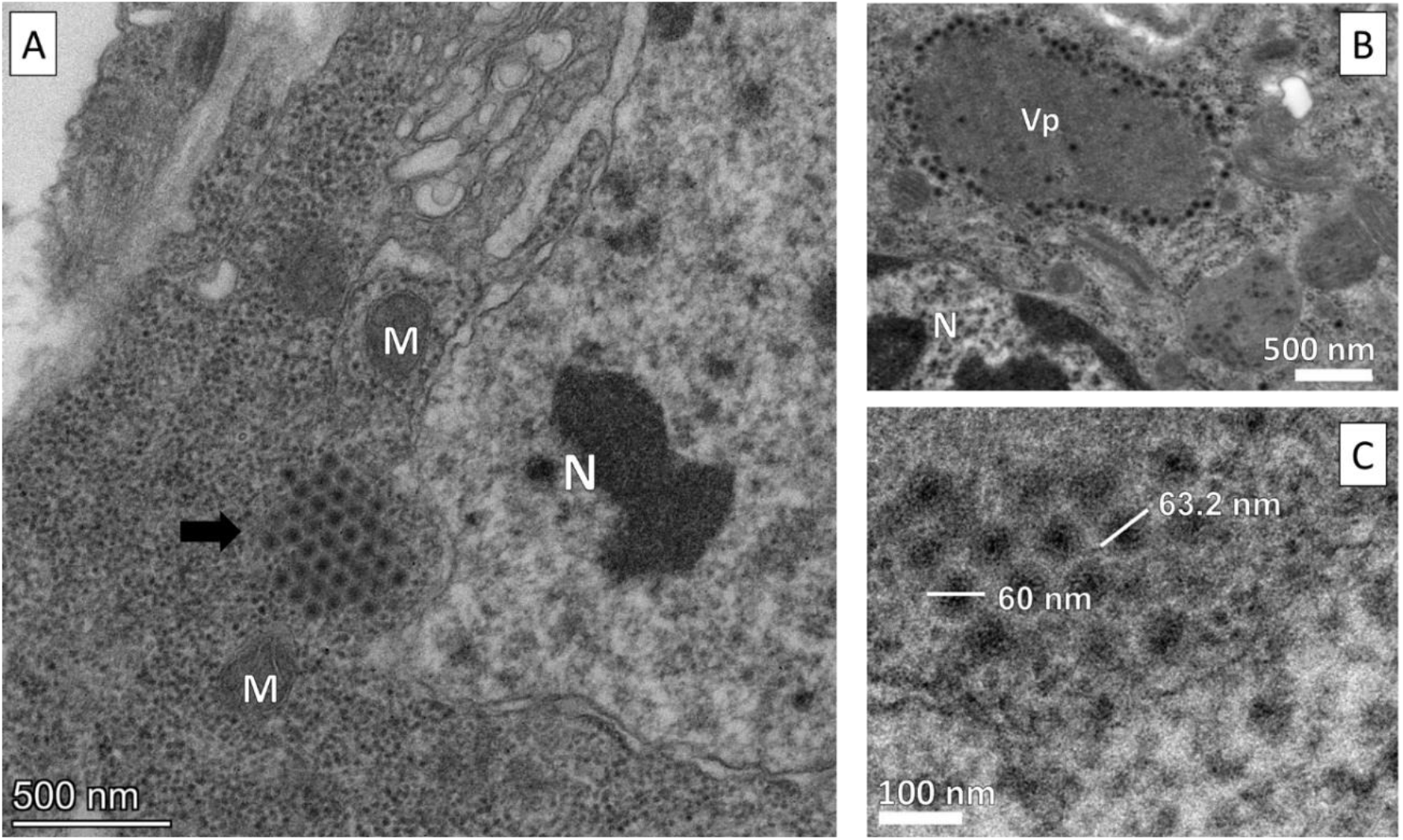
Transmission electron microscope images of AnvRV particles in *A. vladimiri* follicle cells. **A)** a cluster of virus particles (black arrow), **B)** a viroplasm (Vp), **C)** high magnification of a small cluster of virions. N, nucleus; M, mitochondria. Scale bars; A) 500 nm, B) 500nm, C) 100nm.

## Discussion

The viral community of the parasitoid *A. vladimiri*, an agriculturally important biological control agent of mealybugs, is dominated by two positive ssRNA viruses from the order *Picornavirales*, which have sequence similarity to viruses of economically important insects, and a dsRNA segmented genome, assigned to the *Idnoreovirus* genus of the *Reoviridae* family. The latter was localized in the ovaries of the *A. vladimiri*, and was not found in the mealybug hosts, suggesting it specifically infects the wasp.

As most insect viromes are yet to be explored, it was not surprizing to discover new viruses in this host (e.g. Shi et al. 2016). Nevertheless, to the best of our knowledge, other than this study there is only one available detailed report on viruses in wasps of the family Encyrtidae, which reported the genome sequence of one Picorna-like virus in the parasitoid *Diversinervus elegans* (Wu et al., 2021).

Most reoviruses reported to infect insects were found in Dipterans and in Lepidopterans (most of them in species that are important agricultural pests), and are assigned to the genera *Cypovirus* (16 accepted species, which include >75 isolates), and *Idnoreovirus* (7 accepted species, >10 isolates) (Matthijnssens et al., 2022). Some are reported to cause chronic diarrheal disease to larvae, with mild-severe symptoms, while many others have no apparent symptoms on their host (Matthijnssens et al., 2022).

Out of the few reports on reoviruses infecting parasitoids, even fewer have tested for phenotypic effects of the virus on its host. Renault et al., (2005) and Graham et al., (2008) both found no evidence for phenotypic effects of reoviruses inhabiting *Diadromus pulchelus* and *Phobocampe tempestiva* respectively. However, in some cases, reoviruses evidently cause significant effects on their parasitoid host. Renault et al., (2003) reported that during parasitization *Diadromus pulchellus* females inject a reovirus into the lepidopteran host together with the egg, where like the PDV effect, it inhibits the immune response and allows the full development of the parasitoid larva. In contrast, the cypovirus that infects both *Campoletis sonorensis* parasitoid and its lepidopteran host (in which it cannot replicate), seems to be beneficial to the lepidopteran as it reduces percent of mortality from parasitism (Deacutis, 2012).

Viruses of the families *Dicistroviridae* (15 accepted species, which include many more isolates) and *Iflaviridae* (15 accepted species, >80 isolates) are more commonly associated with insects, but similarly to reoviruses, in most cases their phenotype in the insect host is unknown (Valles et al., 2017a, 2017b). Reported effects of viruses from those families are mostly pathogenic (e.g., Acute bee paralysis virus, Drosophila C viruse, Cricket paralysis virus, for *Dicistroviridae*, and Varroa destructor virus1, Deformed wing virus for *Iflaviridae*). Conversely, at least one symbiotic relationship of those viruses with parasitoids was reported. The iflavirus DcPV aids the *Dinocampus coccinellae* to paralyze its ladybeetles host, as it replicates in the ladybeetle brain possibly leading to behaviour changes that protect the developing parasitoid (Dheilly et al., 2015). Another symbiotic relationship was found between a cripavirus (*Dicistroviridae*) and *Drosophila* flies, where the virus seems to increase the fecundity of individuals that carry it (Zhang et al., 2021).

As mentioned above, in most cases the relationship of the virus with its insect host is unknown. As *A. vladimiri* wasps are reared in our lab for many generations, with no apparent disease symptoms, it can be assumed that under the tested rearing conditions (especially the AnvRV, which is fixed in the mass-reared population), the virus is at least not pathogenic, or even may be beneficial to the wasp. Nonetheless, all scenarios are to be taken into consideration when speculating on the relationship of those three novel viruses with *A.vladimiri*, including the possibility that those viruses don’t have any phenotypic effect, and are spread in the population through efficient vertical and or horizontal transmission (e.g. Lüthi et al., 2020; Martinez et al., 2016; Perreau & Moran, 2022). Future study should address these possibilities.

Screening the wasp populations showed that while the two ssRNA viruses are not present in all individuals, the AnvRV is fixed in the lab-reared lines, suggesting either efficient transfer of this virus, whether vertical, horizontal or both (e.g. Varaldi et al., 2006). It may also suggest that the specific rearing conditions favour the carrying of the virus (e.g. Himler et al., 2011; Coffman et al., 2020). However, due to the low prevalence of the AnvRV in field populations of *A. vladimiri*, we conclude that the interaction between the two is not obligatory. Additionally, while the two ssRNA viruses were detected in virion extractions of unparasitized mealybugs, the AnvRV was not, meaning it is a wasp symbiont. On the other hand, the two ssRNA viruses may originate from the mealybug, or they can infect both the wasp and its host (e.g. Zhang et al., 2021)).

The TEM images of virus particles in follicle cells of *A. vladimiri*, resemble the typical pattern found in viruses from the family Reoviridae, which are known to be found in the cytoplasm, to be round shaped, 60-70nm long, and may be occluded in viroplasms (Shah et al., 2017). All species of this genus were detected only in insects (Matthijnssens et al., 2022). Such a localization of a microbe in the ovary, strongly suggests vertical maternal transfer, which in turn may hint on long evolutionary relationships, and perhaps mutualism, between the two (Zchori-Fein and Bourtzis, 2011; Perlmutter and Bordenstein, 2020).

The evolutionary arms race between parasitoids and their hosts have led for several defence mechanisms that hosts use to escape parasitism. In response to invasion by an endo-parasitoid egg (or other foreign organisms), immune haemocytes in mealybugs may aggregate around the inserted egg, and form a melanized capsule, which eliminates its development (Blumberg, 1997). The rates of this encapsulation response to *A. vladimiri* parasitism is known to differ between *P. citri* and *P. ficus* hosts species (Blumberg et al., 1995; Suma et al., 2012). it can be speculated that AnvRV aids the wasp to overcome encapsulation, like DpRV2 does in the *Diadromus pulchellus - Acrolepiopsis assectella* system (Renault et al., 2003), but with different efficiency in different *Planococcus* hosts. If this speculation is indeed true then the different prevalence between lab and field lines can be explained by the fact that the field population originated in vineyards where *P. ficus* is the main host for *A. vladimiri*, whereas the lab population develops in *P. citri* hosts.

Overall results of the screening of all ten segments in *A. vladimiri* specimens, were conclusive and indicate that all segments belong to the same virus. However, four segments failed to amplify in between one-four out of 16 specimens tested. This might suggest that those four segments have lower copy numbers than the other five. The detection of segment number four (k119_19) in two specimens of the negative population can be explained by either technical error, or by contamination.

From the pattern of *Wolbachia* and AnvDV prevalence in the 24 field collected females, where the presence of those two microorganisms is somewhat correlated, it can be speculated that some interaction takes place between those two. Reports on *Wolbachia* interaction with viruses focus on associations with human viral pathogens vectored by insects and insect diseases caused by viruses. For example, *Wolbachia* protects *Drosophila* flies from the Drosophila C virus (Family: *Dicistroviridae*) (e.g. Hedges et al., 2008), and inhibits the Dengue Fever and Chikungunya pathogens in humans (DENV and CHIKV, both also positive ssRNA viruses), (e.g. Hoffmann et al., 2011; Walker et al., 2011). This effect is also detected in natural populations of *Drosophila* and is however virus-specific (Cogni et al., 2021). Those inhibitions were generally correlated with lower viral copy number of the viruses. In other cases, endosymbionts do correlate positively with viral presence and transmission (e.g Gottlieb et al., 2010; Kliot et al., 2014), such as reported here for *A. vladimiri*, where a significant positive correlation was found between the presence of the bacterium and the AnvDV in the field population, and also a higher prevalence in the lab strain infected by *Wolbachia*. This is intriguing since it could suggest a positive effect of *Wolbachia* on virus transmission, may further indicate that this virus and *Wolbachia* interact and clearly needs to be further investigated.

Interestingly, in oppose to most species of Dicistroviridae members, which have a ‘Di-cistronic’ genome (the trait which is the source of the family’s name), only one polypeptide ORF could be identified in AnvDV. This is more similar to other families in the *Picornavirales* order, such as *Iflaviridae* and *Picornaviridae*, however in those families the genes are in opposite order to the order that was found in the AnvDV, where non-structural proteins are closer to the 3’ end (Bonning and Miller, 2010). As far as we know, such genomic arrangement was found only in a few viruses, most of which are from aquatic invertebrates (Shi et al., 2016, see supplementary data #33; Cheng et al., 2021; Wu et al., 2021). This rare finding suggests that this family has a broader diversity than previously known.

## Conclusion

Most studies on viruses in insects focus on pathogenic interactions or transient in vectors to plants or animals. This is one of the first attempts to characterize the virome of a biocontrol agent, that overall seems to be healthy. The discovery of three new RNA viruses in a single population with no apparent diseases, highlights the richness of this hidden community of viruses. This community certainly deserves further investigation in order to clarify its transmission along generations and the phenotypic effects associated. If one (or more) of the three viruses influences the efficiency of *A. vladimiri* as a natural enemy, whether positively or negatively, this knowledge can be applied to improve the biological control of mealybug pests.

## Methods

### Insect rearing and field collections of parasitoids

*Anagyrus vladimiri* wasps were obtained from ‘BioBee Sde Eliyahu Ltd’ mass-rearing (MR) facility (Sde Eliyahu, Israel), and used to establish a *Wolbachia*-positive line (W^+^), from which a *Wolbachia-free* line (W^-^) was established by antibiotic treatments (Izraeli et al., 2020). The two lines were reared under controlled conditions of 26±1°C, 60±20% RH, and 16L:8D photoperiod regime on the citrus mealybugs *Planococcus citri*, feeding on sprouted potatoes *Solanum toberosum*.

Additional *A. vladimiri* individuals were collected in the field during the summer of 2020, using funnel insect traps placed in an unsprayed vineyard in northern Israel (32.7200N, 35.1903E). The traps (n=10), baited with mealybugs-infested potatoes, were retrieved from the field after one week and parasitized mealybugs were incubated in the lab. The *A. vladimiri* adults which emerged (emergence was achieved from three out of the 10 traps, the other seven were empty), were used either for screening viruses’ prevalence (n=24, results in Fig. 3), and others were let to parasitize in the lab, establishing the ‘Field’ line (n=~20 female foundresses). One generation later, eight females of this Field line were individually placed in cups and allowed to oviposit in mealybugs. Mothers and three of their emerged offspring were then tested for the presence of the AnvRV and *Wolbachia*. All the cups in which both symbionts were not detected (n=7, since *Wolbachia* was detected in one mother) were pooled to establish an uninfected *A. vladimiri* line (termed hereafter ‘Field-RV^-^’ line).

The origins of the five wasp lines, and the various experiments they were used for are summarized in Table 1.

### *Enzymatic characterization of* A. vladimiris’ *virome*

To characterize the viral community inhabiting *A. vladimiri*, viral nucleic acids (VNAs) were purified following (Martinez et al., 2016). Briefly, viral capsids were purified from 0.5 g of MR *A. vladimiri* wasps (pool of >1,500 individuals), by filtration, nuclease treatment (RNaseA and DNaseI) and centrifugation. To classify the genome type of potential viruses (RNA/DNA, double/single strand), VNAs were then extracted by SDS, subjected to different nuclease treatments, and then examined on an 0.8% agarose gel (Martinez et al., 2016). A non-treated VNA sample, a dsRNA ladder (Phi6, Invitrogen, MA, USA), and a DNA ladder (1Kb+, Invitrogen, MA, USA) were used as controls.

### Sequencing strategy

To further reveal all viral candidates inhabiting the wasp, VNAs were extracted from purified viral capsids using the All-In-One DNA/RNA Miniprep Kit (Biobasic, ON, Canada). Since the gel images produced in this process showed no indication of viral DNA molecules, only samples with RNA VNAs were further examined and sequenced. Libraries were prepared by the TruSeq RNA Ribo-Zero kit (Illumina, CA, USA) and sequenced by Illumina novaseq 6000 with 100 bp paired-end reads.

Since the sequence yield was very high (~16 million paired-end reads, where suspected viruses’ genome size is up to 30 thousand bp), the data was randomly subsampled to three smaller datasets, including 10^5 paired reads, 10^6 paired reads and 2*10^6 paired reads. The subsampled datasets were *de-novo* assembled using ‘Megahit’ version 3.4 with default parameters. Comparing assembly results of the three subsampled datasets to the original full dataset, revealed that most of the contigs are identical, with a few additional small contigs and the 10^6 paired reads dataset was chosen as the most suitable for further analysis (average fold (sometimes termed ‘depth’) per bp = 4.2*10^4). The resulting contigs were searched for homologous similarity against both the nr and the swissprot databases using BLASTx, targeting viruses, bacteria and Hymenoptera (taxids 10239, 2, 7399 respectively). The following parameters were changed from default to increase sensitivity: matrix – BLOSUM45, Gap costs – 12;2. An additional PSI-Blastp analysis, using the protein sequences predicted by the ORFfinder online tool (RRID:SCR_016643), was used to identify similarity of contigs that Blastx failed to identify.

### Annotation

All contigs were subjected to an open reading frame (ORF) search using ORFfinder online tool (RRID:SCR_016643), with start codon including alternative initiation codons. The length of the un-translated regions (UTR), included the 5’ and 3’UTRs, was calculated as contig length minus the ORF length. As the ORFfinder tool may predict also false proteins, a calculation of UTR divided by the contig length was done to estimate the likeliness of the contig to harbour a ‘true’ protein, the higher the value the lower chance to have a ‘true’ protein.

The UTRs of the ten identified segments of AnvRV were searched for conserved sequence motifs in the 5’ and 3’ end termini. This was done by subjecting those UTR sequences to a BLAST search against themselves, with the automatic adjusted parameters for short sequences of NCBI BLASTn suite.

Further annotation analysis of the viral genomes was done by subjecting all the ORFs to the NCBI batch-CDD tool (Marchler-Bauer et al., 2015), against the CDD v.3.19 database to search for conserved domains. For relevant viruses a search for internal ribosome entry sites (IRES) was done by the IRESite online tool (IRESite: The database of experimentally verified IRES structures, 2022) using the NUC4.4 matrix.

### Phylogeny

To construct phylogenetic trees of the three novel viruses, the RdRP amino acid sequence of each virus was aligned together with ~20 sequences of other viruses from the corresponding family. Representative sequences were chosen from a BLASTp search and from the supplementary material of Shi et al. 2016, to represent the whole family (or subfamily for the AnvRV). As the AnvDV and AnvIFV belong to the same *Picornavirales* order, one tree was constructed for both of them together. Multiple sequence alignment was conducted by Clustal Omega and a maximum likelihood tree was constructed with the LG substitution model using PhyML V.3 (Sievers et al., 2011). Branch support was measured by approximate likelihood ratio tests (SH-aLRT; Guindon et al., 2010). The outgroups chosen for the AnvRV tree were two mammal infecting viruses from the subfamily *Sedoreovirinae*, and a poliovirus sequence was used for the *Picornavirales* tree.

### *Virion purification from* Anagyrus vladimiri *and* Planococcus citri

To test whether any of the three RNA viruses originate from the mealybug hosts, virions were purified from pools of: i) Mass-reared (MR) W^+^ *A. vladimiri* (~0.7 g), ii) Field-collected *A. vladimiri* line without the AnvRV (RV^-^) (~0.2 g), iii) un-parasitiezed *P. citri*,(~0.7 g) and iv) *P. citri*, four days after parasitization by MR *A. vladimiri*, (~0.3 g) (see Table 1 for details). Purification was done according to Luria et al., (2020), with minor modifications. Briefly, the insects were homogenized in a TBE buffer, supernatant was filtered through a 0.45 μm membrane, placed on a 30% sucrose cushion, and subjected to ultracentrifugation at 242,922 g for 2 h. Then, RNA was extracted from the pellet using the Viral RNA extraction kit (Bioneer, Daejeon, South Korea). Detection of the three viruses (AnvRV, AnvIFV and AnvDV) in those samples was carried by cDNA synthesis and PCRs as described below.

### PCR detections of the three viruses

Detection of RNA viruses in *A. vladimiri* individuals was obtained by RNA extraction (RNeazy plus kit, Qiagen, Düsseldorf, Germany) according to manufacturer protocol, with addition of Dithiothreitol to homogenizing buffer, to inhibit degradation by RNase. RNA samples were reverse-transcribed to cDNA using the RT-PCRbio kit (PCR Biosystems, London, UK), with no-template and no-enzyme controls. cDNA was used as templates for diagnostic PCR to identify presence/absence of the three viruses. Specific primers were designed for each virus and each viral genomic segment according to the contig sequences identified from the RNA-seq data (Supp. table 1).

### Transmission electron microscope (TEM)

*Anagyrus vladimiri* females from the W^+^RV^+^ line (see Table 1 for details) were used to test for the presence of viruses in the reproductive tissues of the parasitoids. Dissected *A. vladimri* ovaries were fixed overnight with 1% Paraformaldehyde, 2.5% Glutaraldehyde in 0.1 M Cacodylate buffer pH 7.4 containing 5 mM CaCl2, washed in 0.1 M cacodylate, post fixed for 1 hour in with 1% OsO4 and 5 mM CaCl2 in Cacodylate buffer. Then the samples went through dehydration in ethanol and embedded in epon 812 (EMS). The blocks were cut with a diamond knife using a UC7 ultramicrotome (Leica,Wetzlar, Germany). Sections of ~75nm were collected on grids, contrast stained for 10 min with 2% uranyl acetate and visualized in a TEM (L120C, Talos, Thermo Fisher Scientific, MA, USA).

## Data availability

Accession numbers to be received.

## Acknowledgements

The authors would like to thank Prof. A. Dombrovsky for useful advice, E. Erel for insect rearing consultancy and Dr. L. Shaulov for preparation and visualization of ovary slides in TEM.

## References

Avidov, Z., Rosseler, Y., and Rosen, D. (1967). Studies on an Israel strain of *Anagyrus pseudococci* (Girault) [Hym., Encyrtidae]. II. Some biological aspects. Entomophaga 12, 111–118. doi: 10.1007/BF02370607.

Blumberg, D. (1997). Parasitoid encapsulation as a defense mechanism in the Coccoidea (Homoptera) and its importance in biological control. Biological Control 8, 225–236. doi:10.1006/BCON.1997.0502.

Blumberg, D., Klein, M., and Mendel, Z. (1995). Response by encapsulation of four mealybug species (Homoptera: Pseudococcidae) to parasitization by *Anagyrus pseudococci*. Phytoparasitica 23, 157–163. doi:10.1007/BF02980975.

Bonning, B. C., and Miller, W. A. (2010). Dicistroviruses. Annual Review of Entomology 55, 129–150. doi:10.1146/ANNUREV-ENTO-112408-085457.

Bugila, A. A. A., Franco, J. C., da Silva, E. B., and Branco, M. (2015). Suitability of five mealybug species (Hemiptera, Pseudococcidae) as hosts for the solitary parasitoid *Anagyrus sp. nr. pseudococci* (Girault) (Hymenoptera: Encyrtidae). Biocontrol Science and Technology 25, 108–120. doi:10.1080/09583157.2014.952711.

Wu, C. Y., Zhao, Y. J., and Zhu, J. Y. (2021). Genome sequence of a novel member of the order *Picornavirales* from the endoparasitoid wasp *Diversinervus elegans*. Archives of Virology 166, 295–297. doi: 10.1007/s00705-020-04824-y.

Cheng, R. L., Li, X. F., and Zhang, C. X. (2021). Novel Dicistroviruses in an unexpected wide range of invertebrates. Food and Environmental Virology 13, 423–431. doi:10.1007/s12560-021-09472-2.

Coffman, K. A., Harrell, T. C., and Burke, G. R. (2020). A mutualistic Poxvirus exhibits convergent evolution with other heritable viruses in parasitoid wasps. Journal of Virology 94, e02059–19. doi:10.1128/JVI.02059-19.

Cogni, R., Ding, S. D., Pimentel, A. C., Day, J. P., and Jiggins, F. M. (2021). *Wolbachia* reduces virus infection in a natural population of *Drosophila*. Communications Biology 4, 1327. doi:10.1038/s42003-021-02838-z.

Deacutis, J. (2012). The characterization and biological effects of a novel cypovirus on the Heliothis virescens and Campoletis sonorensis host-parasitoid system. University of Kentucky.

Dheilly, N. M., Maure, F., Ravallec, M., Galinier, R., Doyon, J., Duval, D., et al. (2015). Who is the puppet master? Replication of a parasitic wasp-associated virus correlates with host behaviour manipulation. Proceedings of the Royal Society B: Biological Sciences 282, 20142773. doi:10.1098/rspb.2014.2773.

Drew, G. C., Stevens, E. J., and King, K. C. (2021). Microbial evolution and transitions along the parasite–mutualist continuum. Nature Reviews Microbiology 19, 623–638. doi:10.1038/s41579-021-00550-7.

Godfray, H. J. C. (1994). Parasitoids: Behavioral and Evolutionary Ecology. Princeton University Press.

Gottlieb, Y., Zchori-Fein, E., Mozes-Daube, N., Kontsedalov, S., Skaljac, M., Brumin, M., et al. (2010). The transmission efficiency of tomato yellow leaf curl virus by the whitefly *Bemisia tabaci* is correlated with the presence of a specific symbiotic bacterium species. Journal of Virology 84, 9310–9317. doi: 10.1128/jvi.00423-10.

Graham, R. I., Rao, S., Sait, S. M., Attoui, H., Mertens, P. P. C., Hails, R. S., et al. (2008). Sequence analysis of a reovirus isolated from the winter moth Operophtera brumata (Lepidoptera: Geometridae) and its parasitoid wasp *Phobocampe tempestiva* (Hymenoptera: Ichneumonidae). Virus Research 135, 42–47. doi:10.1016/j.virusres.2008.02.005.

Guindon, S., Dufayard, J. F., Lefort, V., Anisimova, M., Hordijk, W., and Gascuel, O. (2010). New algorithms and methods to estimate maximum-likelihood phylogenies: Assessing the performance of PhyML 3.0. Systematic Biology 59, 307–321. doi:10.1093/SYSBIO/SYQ010.

Hedges, L. M., Brownlie, J. C., O’Neill, S. L., and Johnson, K. N. (2008). *Wolbachia* and virus protection in insects. Science 322, 702–702. doi:10.1126/science.1162418.

Herniou, E. A., Huguet, E., Thézé, J., Bézier, A., Periquet, G., and Drezen, J.-M. (2013). When parasitic wasps hijacked viruses: genomic and functional evolution of polydnaviruses. Philosophical Transactions of the Royal Society B: Biological Sciences 368, 20130051. doi:10.1098/RSTB.2013.0051.

Himler, A. G., Adachi-Hagimori, T., Bergen, J. E., Kozuch, A., Kelly, S. E., Tabashnik, B. E., et al. (2011). Rapid spread of a bacterial symbiont in an invasive whitefly is driven by fitness benefits and female bias. Science 332, 254–256. doi:10.1126/science.1199410.

Hoffmann, A. A., Montgomery, B. L., Popovici, J., Iturbe-Ormaetxe, I., Johnson, P. H., Muzzi, F., et al. (2011). Successful establishment of *Wolbachia* in *Aedes* populations to suppress dengue transmission. Nature 476, 454–457. doi:10.1038/nature10356.

IRESite: The database of experimentally verified IRES structures (2022). http://www.iresite.org/IRESite_web.php?page=blastsearch&search_type=blast_iress [Accessed January 30, 2022].

Izraeli, Y., Lalzar, M., Mozes-Daube, N., Steinberg, S., Chiel, E., and Zchori-Fein, E. (2020). *Wolbachia* influence on the fitness of *Anagyrus vladimiri* (Hymenoptera: Encyrtidae), a bio-control agent of mealybugs. Pest Management Science 77, 1023–1034. doi:10.1002/ps.6117.

Khramtsov, N. v., Woods, K. M., Nesterenko, M. v., Dykstra, C. C., and Upton, S. J. (1997). Virus-like, double-stranded RNAs in the parasitic protozoan *Cryptosporidium parvum*. Molecular Microbiology 26, 289–300. doi:10.1046/j.1365-2958.1997.5721933.x.

Kliot, A., Cilia, M., Czosnek, H., and Ghanim, M. (2014). Implication of the bacterial endosymbiont *Rickettsia* spp. in interactions of the whitefly *Bemisia tabaci* with tomato yellow leaf curl virus. Journal of Virology 88, 5652–5660. doi: 10.1128/jvi.00071-14.

Luria, N., Smith, E., Lachman, O., Laskar, O., Sela, N., and Dombrovsky, A. (2020). Isolation and characterization of a novel cripavirus, the first Dicistroviridae family member infecting the cotton mealybug *Phenacoccus solenopsis*. Archives of Virology 165, 1987–1994. doi:10.1007/s00705-020-04702-7.

Lüthi, M. N., Vorburger, C., and Dennis, A. B. (2020). A Novel RNA Virus in the parasitoid wasp *Lysiphlebus fabarum*: Genomic structure, prevalence, and transmission. Viruses 12, 59. doi: 10.3390/V12010059.

Marchler-Bauer, A., Derbyshire, M. K., Gonzales, N. R., Lu, S., Chitsaz, F., Geer, L. Y., et al. (2015). CDD: NCBI’s conserved domain database. Nucleic Acids Research 43, D222–D226. doi:10.1093/NAR/GKU1221.

Martinez, J., Lepetit, D., Ravallec, M., Fleury, F., and Varaldi, J. (2016). Additional heritable virus in the parasitic wasp *Leptopilina boulardi*: Prevalence, transmission and phenotypic effects. Journal of General Virology 97, 523–535. doi:10.1099/jgv.0.000360.

Matthijnssens, J., Attoui, H., Bányai, K., Brussaard, C. P. D., Danthi, P., Vas, M., et al. (2022). ICTV Virus Taxonomy Profile: Reoviridae. Journal of General Virology.

Perlmutter, J. I., and Bordenstein, S. R. (2020). Microorganisms in the reproductive tissues of arthropods. Nature Reviews Microbiology 18, 97–111. doi:10.1038/s41579-019-0309-z.

Perreau, J., and Moran, N. A. (2022). Genetic innovations in animal–microbe symbioses. Nature Reviews Genetics 23, 23–39. doi: 10.1038/s41576-021-00395-z.

Renault, S., Bigot, S., Lemesle, M., Sizaret, P.-Y. Y., and Bigot, Y. (2003). The cypovirus Diadromus pulchellus RV-2 is sporadically associated with the endoparasitoid wasp *D. pulchellus* and modulates the defence mechanisms of pupae of the parasitized leek-month, *Acrolepiopsis assectella*. Journal of General Virology 84, 1799–1807. doi:10.1099/vir.0.19038-0.

Renault, S., Stasiak, K., Federici, B., and Bigot, Y. (2005). Commensal and mutualistic relationships of reoviruses with their parasitoid wasp hosts. Journal of Insect Physiology 51, 137–148. doi:10.1016/j.jinsphys.2004.08.002.

Shah, P. N. M., Stanifer, M. L., Höhn, K., Engel, U., Haselmann, U., Bartenschlager, R., et al. (2017). Genome packaging of reovirus is mediated by the scaffolding property of the microtubule network. Cellular Microbiology 19, e12765. doi:10.1111/cmi.12765.

Shi, M., Lin, X. D., Tian, J. H., Chen, L. J., Chen, X., Li, C. X., et al. (2016). Redefining the invertebrate RNA virosphere. Nature 540, 539–543. doi:10.1038/nature20167.

Sievers, F., Wilm, A., Dineen, D., Gibson, T. J., Karplus, K., Li, W., et al. (2011). Fast, scalable generation of high-quality protein multiple sequence alignments using Clustal Omega. Molecular Systems Biology 7, 539. doi:10.1038/MSB.2011.75.

Sorrentino, S., Carsana, A., Furia, A., Doskocil, J., and Libonati, M. (1980). Ionic control of enzymic degradation of double-stranded RNA. Biochimica et Biophysica Acta 609, 40–52. doi:10.1016/0005-2787(80)90199-9.

Suma, P., Mansour, R., Russo, A., La Torre, I., Bugila, A. A. A., and Franco, J. C. (2012). Encapsulation rates of the parasitoid *Anagyrus* sp. nr. pseudococci, by three mealybug species (Hemiptera: Pseudococcidae). Phytoparasitica 40, 11–16. doi:10.1007/s12600-011-0199-8.

Valles, S. M., Chen, Y., Firth, A. E., Guérin, D. M. A., Hashimoto, Y., Herrero, S., et al. (2017a). ICTV virus taxonomy profile: Dicistroviridae. Journal of General Virology 98, 355–356. doi:10.1099/JGV.0.000756.

Valles, S. M., Chen, Y., Firth, A. E., Guérin, D. M. A., Hashimoto, Y., Herrero, S., et al. (2017b). ICTV virus taxonomy profile: Iflaviridae. Journal of General Virology 98, 527–528. doi:10.1099/JGV.0.000757/CITE/REFWORKS.

Varaldi, J., Boulétreau, M., and Fleury, F. (2005). Cost induced by viral particles manipulating superparasitism behaviour in the parasitoid *Leptopilina boulardi*. Parasitology 131, 161–168. doi:10.1017/S0031182005007602.

Varaldi, J., Fouillet, P., Ravallec, M., López-Ferber, M., Boulétreau, M., and Fleury, F. (2003). Infectious behavior in a parasitoid. Science 302, 1930. doi:10.1126/science.1088798.

Varaldi, J., Gandon, S., Rivero, A., Patot, S., Fleury, F., Gandon, S., et al. (2006). “A newly discovered virus manipulates superparasitism behavior in a parasitoid wasp,” in Insect Symbiosis, ed. Bourtzis, K., and Miller, T. A. Taylor and Francis, (Boca Raton, FL: CRC Press), 141–162. doi:10.1201/9781420005936-11.

Walker, T., Johnson, P. H., Moreira, L. A., Iturbe-Ormaetxe, I., Frentiu, F. D., McMeniman, C. J., et al. (2011). The wMel *Wolbachia* strain blocks dengue and invades caged *Aedes aegypti* populations. Nature 476, 450–453. doi:10.1038/NATURE10355.

Webster, C. L., Longdon, B., Lewis, S. H., and Obbard, D. J. (2016). Twenty-Five new viruses associated with the Drosophilidae (Diptera). Evolutionary Bioinformatics 12, 13–25. doi:10.4137/EBO.S39454.

Wu, H., Pang, R., Cheng, T., Xue, L., Zeng, H., Lei, T., et al. (2020). Abundant and diverse RNA viruses in insects revealed by RNA-Seq analysis: Ecological and evolutionary implications. mSystems 5, 1–14. doi:10.1128/msystems.00039-20.

Zchori-Fein, E., and Bourtzis, K. (2011). Manipulative Tenants; Bacteria Associated with Arthropods. Boca Raton: CRC Press. doi:10.1201/b11008.

Zhang, J., Wang, F., Yuan, B., Yang, L., Yang, Y., Fang, Q., et al. (2021). A novel cripavirus of an ectoparasitoid wasp increases pupal duration and fecundity of the wasp’s *Drosophila melanogaster* host. The ISME Journal 15, 3239–3257. doi:10.1038/s41396-021-01005-w.

